# 10x Genomics Gene Expression Flex is a powerful tool for single-cell transcriptomics of xenograft models

**DOI:** 10.1101/2024.01.25.577066

**Authors:** Oriol Llora-Batlle, Anca Farcas, Doreth Fransen, Nicolas Floc’h, Sara Talbot, Alix Schwiening, Laura Bojko, John Calver, Natasa Josipovic, Kanstantsin Lashuk, Julia Schueler, Andrei Prodan, Dylan Mooijman, Ultan McDermott, PERSIST-SEQ Consortium

## Abstract

The 10x Genomics Gene Expression Flex protocol allows profiling of fixed or frozen material, greatly simplifying the logistics of sample collection, storage and transfer prior to single -cell sequencing. The method makes single-cell transcriptomics possible for existing fresh-frozen or FFPE tissue samples, but also facilitates the logistics of the sampling process, allowing instant preservation of samples. The technology relies on species-specific probes available for human and mouse. Nevertheless, processing of patient-derived (PDX) or cell line (CDX) xenografts, which contain mixed human and mouse cells, is currently not supported by this protocol due to the high degree of homology between the probe sets. Here we show that it is feasible to simultaneously profile populations containing both human and mouse cells by mixing the transcriptome probe sets of both species. *Cellranger* outputs a count table for each of the species allowing evaluation of the performance of the different probe sets. Cross-reactive probes are greatly outperformed by the specific probe hybridizations leading to a clear difference in the recovery of UMIs and unique genes per cell. Furthermore, we developed a pipeline that removes cross-reactive signal from the data and provides species-specific count tables for further downstream analysis. Hence, the 10x Genomics Gene Expression Flex protocol can be used to process xenograft samples without the need for separation of human and mouse cells by flow sorting and allows analysis of the human and mouse single-cell transcriptome from each sample. We anticipate it will be increasingly used for single-cell sequencing of cancer cell line and patient-derived xenografts, facilitating the preservation of the samples and allowing the interrogation of both the (human) xenograft and the (mouse) tumor microenvironment at single-cell resolution.

## INTRODUCTION

In 2022, 10x Genomics introduced RNA Flex (formerly known as Fixed RNA Profiling), a probe based single-cell RNA detection method compatible with human and mouse cells. Probe based gene expression analysis dates back to 1977, with the development of Northern blotting, where radioactively labeled DNA probes are used to detect transcripts on a blot^1^. 15 years later, Jim Eberwine used a combination of *in vitro* transcription and Northern blotting to perform the first single-cell RNA detection experiment^2^. It was not until the introduction of microarray technology that probe-based high throughput gene expression analysis became commonplace^3^ and allowed for single-cell RNA gene expression profiling^4^. Second generation sequencing technologies largely made probe-based gene expression analysis obsolete, except for Padlock probe technologies which still allowed for cost effective SNP detection^5^.

In recent years, the field of spatial biology has revisited the use of probe-based methods building from technologies such as single-molecule FISH (smFISH)^6^ and combinatorial barcoding^7^. By coupling the probes to different colors, the spatial location of the RNA of interest can be determined, nevertheless these are limited in multiplexing and quantification capabilities. To circumvent these limitations, modern spatial transcriptomic techniques rely on evaluating tissue samples in a sequential manner after cycles of reagent incubations (i.e. SeqFISH^8^, MERFISH^9^, Xenium^10^) or rely on highly multiplexed approaches that profile regions of interest simultaneously and can be independent of fluorescently labelled probes (i.e. GeoMx^11^, Visium for FFPE samples)^12^. In general, probe-based methods allow for a more sensitive and specific analysis but have limited coverage to the transcripts bound by the probes. Nevertheless, probe-based methods have expanded the capacity to efficiently profile tissue samples preserved as formalin-fixed paraffin-embedded (FFPE) or formalin-fixed (FF). Single-cell methods covering FFPE, FF and snap frozen tissues represented an unmet need for the field until very recently. The 10x Gene Expression Flex kit is one solution to obtain single-cell transcriptomic data from these types of samples^13^.

Tissue samples are usually dissociated and cryopreserved as single-cell suspensions or snap-frozen for nuclei extraction. Tissue dissociation can have a detrimental effect on cell viability and also induce stress responses with resulting changes in gene expression^14^. Additionally, cryopreservation also requires a thawing procedure that usually leads to some cell death (Fig. 1A). Therefore, together with the dissociation stress, samples usually require extra sample processing such as dead cell removal solutions, which extends the time to which cells are exposed^15^. On the other hand, nuclei extraction circumvents some of these hurdles, but only recovers nuclear RNA, which leads to a lower number of genes recovered^16^. With the new Gene Expression Flex kit, tissue samples can be fixed on-site or snap frozen for later processing, therefore preventing sample degradation and solving the logistical challenges often associated with clinical settings; including the inactivation of pathogens present in the samples (Fig 1B).

**Figure 1.**
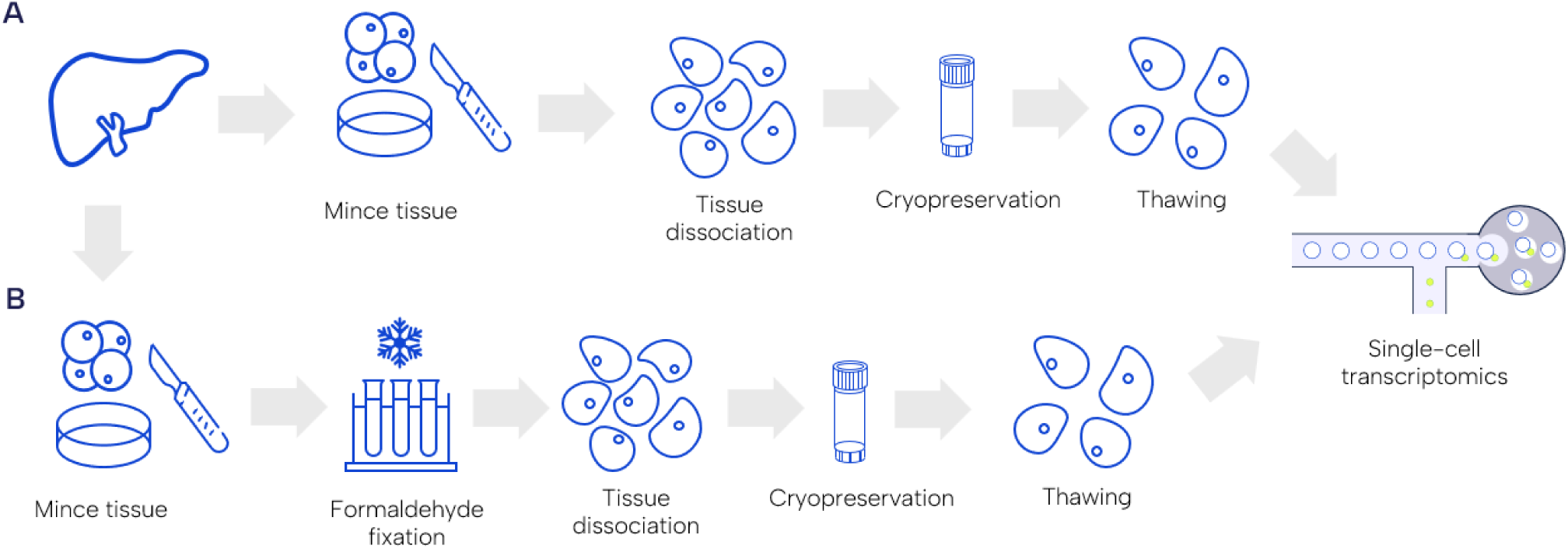
Workflow for processing tissue samples with the 10x Genomics Gene Expression Flex kit compared to the standard pipelines. **A.** Stepwise processing of PDX samples for single-cell transcriptomics. After sampling the tissue from the mouse, the sample is minced into fine pieces to facilitate the enzymatic dissociation with a cocktail of enzymes and obtain a single-cell suspension. If the suspension is of good quality, the cells can be cryopreserved to be shipped. Before processing, cells are thawed and evaluated before proceeding to the single-cell encapsulation and profiling. **B**. When using the Flex kit, minced tissue can be fixed directly, preserving the sample straight away and preventing further cell stress during dissociation, cryopreservation, and thawing.

The 10x Gene Expression Flex kit can be used with either human or mouse whole transcriptome probe sets and the barcoded probes allow for sample multiplexing. Each probe hybridization can target 8,000-10,000 cells and up to 16 samples can be pooled in a single encapsulation reaction in the Chromium X, making the Flex protocol more cost-effective than Single-cell 3’ v3.1 sequencing. Officially the use of human and mouse probe sets in samples containing a mixture of the two species is not supported given the high degree of homology at the transcript level, increasing the chances of probe cross-reactivity. Therefore, despite the advantages in sample preservation and the logistics provided by the Flex kit, PDX models are, in theory, not suitable for this kit.

PDX and CDX consist of the engraftment of human tumoral tissue into immunocompromised mice where the engrafted tissue is sustained by the host and will further develop, simulating the progression and evolution of the tumor. These xenograft models have become widely used in preclinical studies for the identification of biomarkers or testing of new drugs. In PDX models, following engraftment, the human tumor microenvironment (TME) is eventually replaced by the murine host cells. This leads to samples containing a mixture of both human and murine cells. The TME plays a key role in the growth and progression of the tumor^17^, but also has an impact on the sensitivity of the tumors to drugs^18^. Despite the slow replacement of the human stromal cells, there is evidence that the tumor can educate the murine cells to promote its development^19^. Therefore, characterizing the murine TME during therapeutic interventions may be key to understanding the response of the xenograft to the treatment^17^.

To address this question, we tested the feasibility of using the Flex kit to profile samples containing both human and mouse cells and devised the best approach to profile PDX or CDX models with varying degrees of mouse cell infiltration. These situations can be common in models treated with oncology drugs where the tumors shrink and most cells in the sample belong to the mouse host. The barcoding of the probes allows multiplexing samples but also provides a lot of flexibility on the number of cells one can target per sample by altering the representation of each of the samples in the pool.

Here we demonstrate the suitability of the 10x Genomics Gene Expression Flex kit in the profiling of xenograft models and characterizing both the tumor and the TME. The capacity to directly preserve tissue samples has greatly improved sample handling and shipment to the processing laboratories. We foresee that protocols such as the Flex offer the opportunity to simplify the sample collection process for single-cell sequencing, minimize any potential detrimental effects of cell dissociation and cryopreservation and additionally allow the analysis of human and mouse cells from the same sample. Of note, the purpose of this study was not to exhaustively benchmark this protocol against other single-cell fixation processes, and it is likely that similar outcomes can be achieved using other similar commercial applications.

## RESULTS

### The Gene Expression Flex kit can profile human cells in samples containing both human and mouse cells

We first tested whether we could reliably profile human cells in samples containing both human and mouse cells. To do so, we processed cultured PC9 human lung cancer cells spiked with 25% or 50% NIH-3T3 mouse cells using the 10x Genomics Gene Expression Flex kit. The count tables obtained were evaluated using Seurat v4.3^20^, which revealed a clear bi-modal distribution in number of genes and UMI counts per cell in the spiked samples (Fig. 2A). After normalization, highly variable gene selection, scaling and dimensionality reduction, we clustered the cells, obtaining 2 main regions where cells cluster in the spiked samples (Fig. 2B). The clustering reflects the presence of cells with a low number of UMI counts (Fig. 2C). Therefore, we hypothesized that the human probes bind to mouse genes with much lower efficiency and are recovered during the library preparation, resulting in low UMI counts.

**Figure 2.**
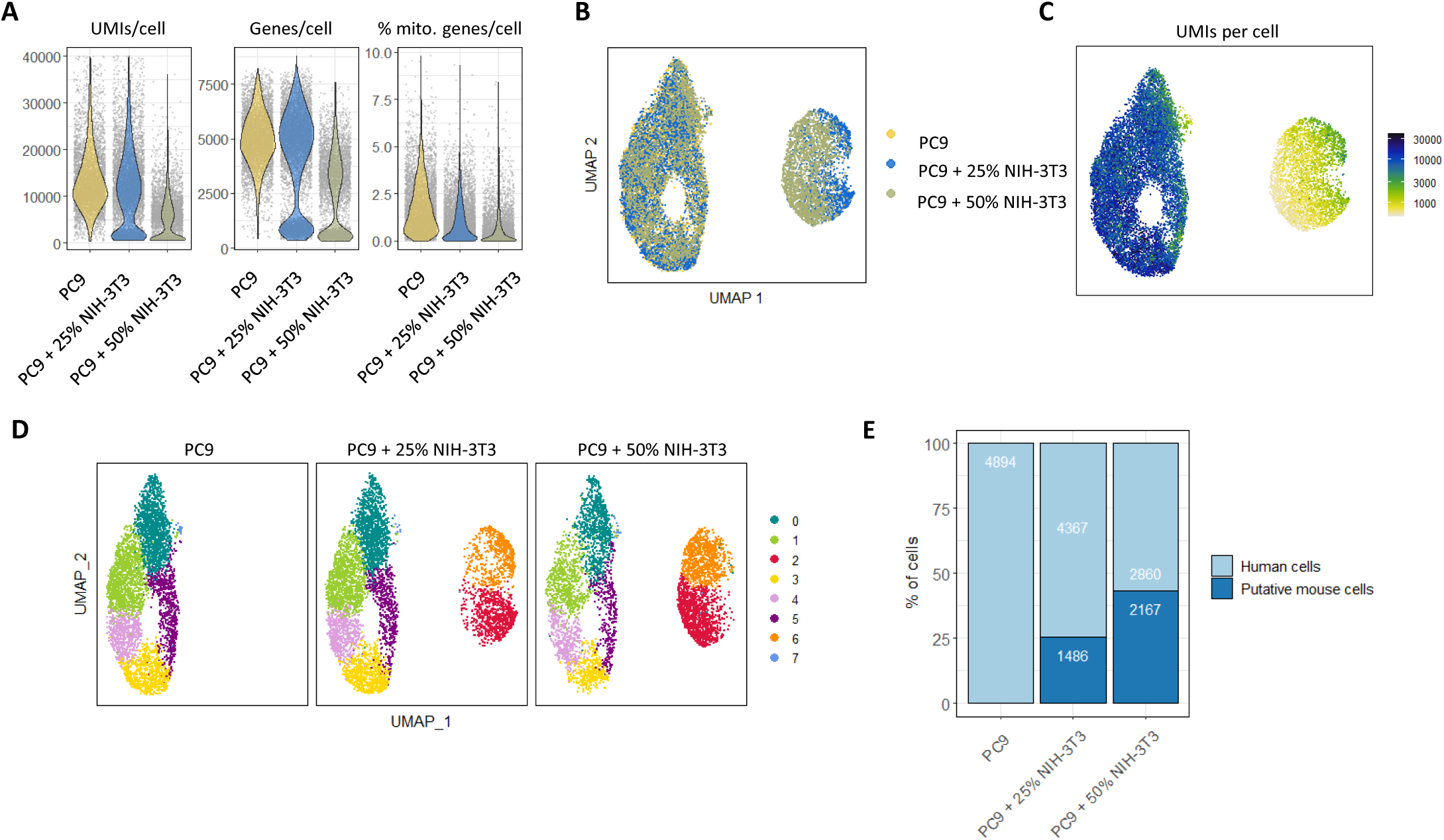
Gene Expression Flex data on mixed *in vitro* samples. **A.** Violin plots showing the distribution of genes, UMI counts and percentage of mitochondrial genes per cell in PC9 (yellow), PC9 spiked with 25% mouse cells (blue) and PC9 spiked with 50% mouse cells (olive). **B.** UMAP visualization of the clustering of the 3 samples. **C.** UMAP visualization with the UMI counts per cell. **D.** UMAP visualization split by sample of origin. **E.** Percentage of human (light blue) and putative mouse (dark blue) cells present in each of the samples when assigning the cells present in the low UMI count clusters as mouse cells. Absolute numbers are shown on top.

We noticed that the number of cells present in the clusters with low UMI counts increased according to the number of mouse cells in the sample (Fig. 2D). In fact, the percentage of cells present within the low UMI clusters matches the number of spiked mouse cells in the sample, with 25.39% and 43.11% of cells for the 25% and 50% spiked samples respectively (Fig. 2E). Therefore, mouse cells are recovered when using human probes but can be easily identified based on the number of UMI counts and genes recovered. This likely reflects the lower efficiency of the cross-reactive probes.

We also asked whether the signal recovered from the mouse cells was due to a subset of probes or not. When looking at the average gene counts in the putative mouse cells cluster, there are around 200 genes with average counts higher than 1, with most of the probes otherwise providing spurious counts (Fig. S1A, Suppl. Table 1).

Next, we tested whether the presence of mouse cells has an impact on the profiling of the human cells. To do so, we compared the average gene counts of the PC9 cells to the ones present in spiked samples, which showed a good correlation between them. Only 15 genes show a significant difference between conditions and most are mitochondrial genes (Fig. S1B-E). When looking at the expression profile of the differentially expressed genes among samples and clusters, no clear pattern emerges, which suggests that none of the clusters is driven by these gene expression differences (Fig. S1E). Additionally, most of the genes showing differential expression in the human cells are very lowly expressed in the putative mouse clusters and the expression levels also correlate with the amount of mouse cells present (Fig. S1F-G). A plausible explanation for the lower gene expression levels could be the competition between human and mouse transcripts for the binding of the probes, leading to lower counts. In fact, mitochondrial genes and some of the other genes detected are mostly profiled by a single probe, which could make the effect of competition more noticeable.

We also assessed whether the reads obtained from the human probes would map to the mouse genome. When using a mouse reference genome, less than 3% of the reads map to the probe set. Overall, the profiling of human cells with the 10x Genomics Flex kit in mixed populations is feasible and provides good quality data.

### Probe mixing and sub-pooling strategies are instrumental to profile both human and mouse cells in mixed samples

We next tested whether profiling of both human and mouse cells within a mixed population is feasible. To do so, we devised 3 strategies (Fig. 3A): The first option is to mix species’ probes tagged with the same barcode; the rationale being that probe mixing is not supported by 10x Genomics and cellranger might not work properly when a cell contains reads from two different probes with distinct barcodes. The second option is to mix species probes with distinct barcodes in the hope that cellranger would work without issues as it should interpret this as having two different samples thereby allowing us to easily identify the origin of the reads. Finally, the third option is to split the sample in two and perform separate hybridizations with the human and mouse probes. This option would provide a reference to which compare the results of option 1 and 2.

**Figure 3.**
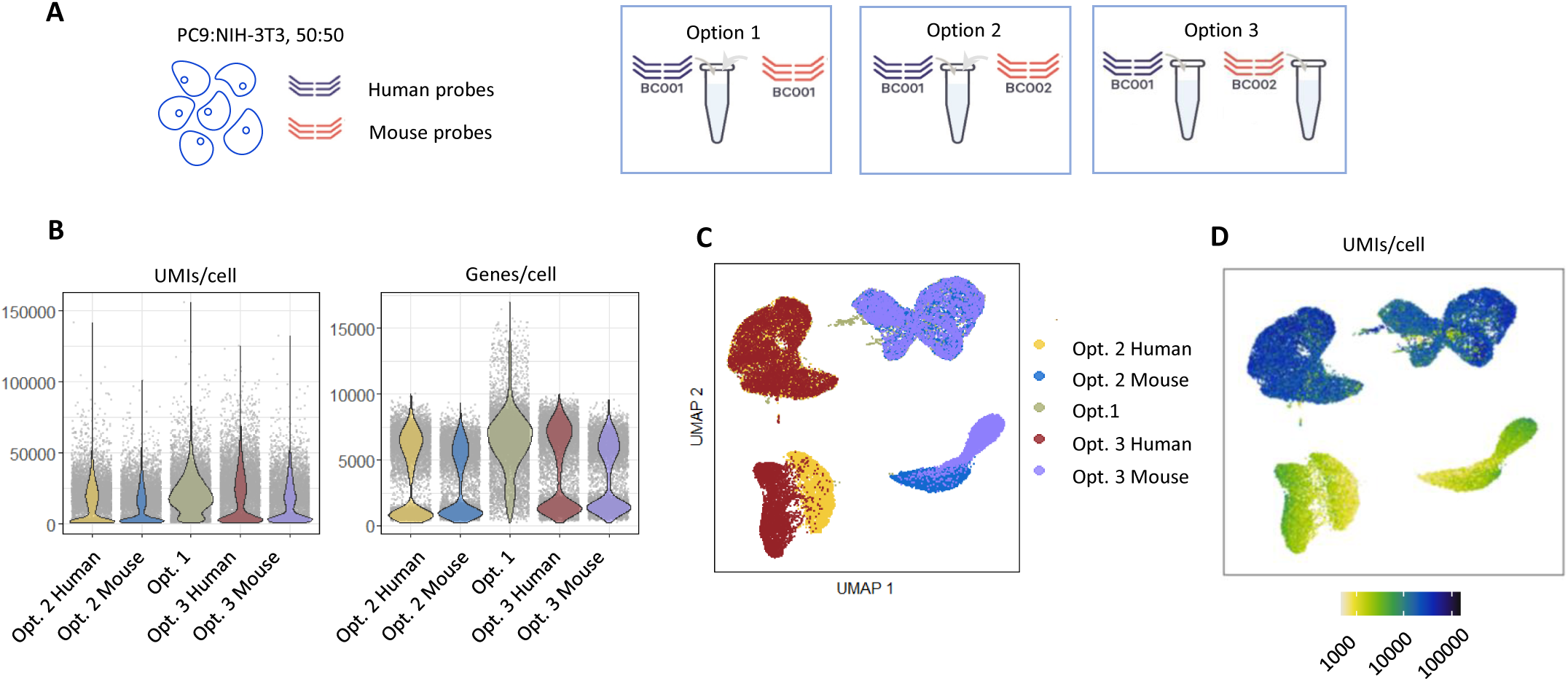
Testing of alternative strategies to profile both human and mouse cells within a mixed sample. **A.** Schematic of the options to profile both human and mouse cells from a mixed sample. Option 1 mixes species probes with the same probe barcode. Option 2 mixes species probes with different probe barcodes. Option 3 splits the sample in two and uses a single species probe set for each hybridization. **B.** Violin plots showing the distribution of genes, UMI counts per cell in option 2 human probes (yellow), option 2 mouse probes (blue), option 1 probes (olive), option 3 human probes (red) and option 3 mouse probes (purple). **C.** UMAP visualization of the clustering obtained from the alternative strategies. **D.** UMAP visualization with the UMI counts per cell of all options.

To test the different options, we first profiled cell cultures containing 50% human and mouse cells (PC9 and NIH-3T3). For de-multiplexing and mapping of the reads, cellranger was set to use a combined human and mouse reference genome and a combined human and mouse probe set. For samples hybridized with different barcodes, the input command was set as if we did 2 hybridizations for the same sample. This allowed us to obtain an individual output for each of the barcodes and to evaluate the contribution of each of the species’ probes independently. Cellranger worked without issues for all the strategies tested. It just provided a warning for samples profiled with the option 2 strategy, given the presence of two barcodes within the same sample.

By inspecting the number of genes and counts per cell, we observed again a bimodal distribution, which is absent in the samples where the same barcode was used (option 1) (Fig. 3B). The clustering revealed again the presence of cells with very low UMI counts that cluster apart in options 2 and 3 (Fig. 3C-D). Therefore, mixing probes with the same barcode is not a suitable option as the cells profiled by the cross-reactive probes cannot be easily identified.

Some mouse probes also cross-react to human transcripts, as shown by the presence of low UMI count clusters in the samples where mouse probes were used. The clustering also shows a perfect overlap of human and mouse cells profiled with the different strategies, respectively (Fig. 3C-D). This suggests that the profiling of the human and mouse cells is not affected by the strategy of choice. In fact, the clustering of the PC9 cells profiled in this experiment is driven by the same genes as in the original sample with only PC9 cells, shown by the top 10 genes expressed per cluster (Fig. S2A-C). Additionally, only 6 genes show significant expression changes between the PC9 cells profiled in these experiments and the PC9 cells alone (Fig. S2D-E), with a good correlation between the average gene counts among samples (Fig. S2F-H). Therefore, both probe mixing with distinct barcodes (option 2) and sub-pooling (option 3) are reliable strategies to profile both human and mouse cells, without severely impacting the profiling of each of the species.

### Cells profiled by the cross-reactive probes are also correctly profiled by the species’ specific probes

One could argue that the cells showing low UMI counts and low number of genes are sub-optimally profiled cells due to the cross-reactivity of the probes, rather than cells corresponding to the other species, or are a product of cross-reactivity artifacts.

To test this, we focused on the sample where we mixed the species probes with different barcodes (option 2). Cellranger profiled this sample as if it consisted of two different hybridizations and therefore produced a separate count table for each of the species’ probes. Any cell present in this sample can interact with only one species probe or with both of them. If this happens, during the single-cell encapsulation, the GEM barcode identifying that single-cell will be incorporated into both the human and mouse probes present in the cell and therefore both the human and mouse count table should contain the same GEM barcode (Fig. 4A). Indeed, in this experiment, most of the cells were present in both the human and mouse count table (Fig. 4B). This means that the cells profiled by cross-reactive probes, which contain low UMI counts in one of the count tables, are also profiled by the correct species probes and can be recovered in the other species’ count table (Fig. 4C-D).

**Figure 4.**
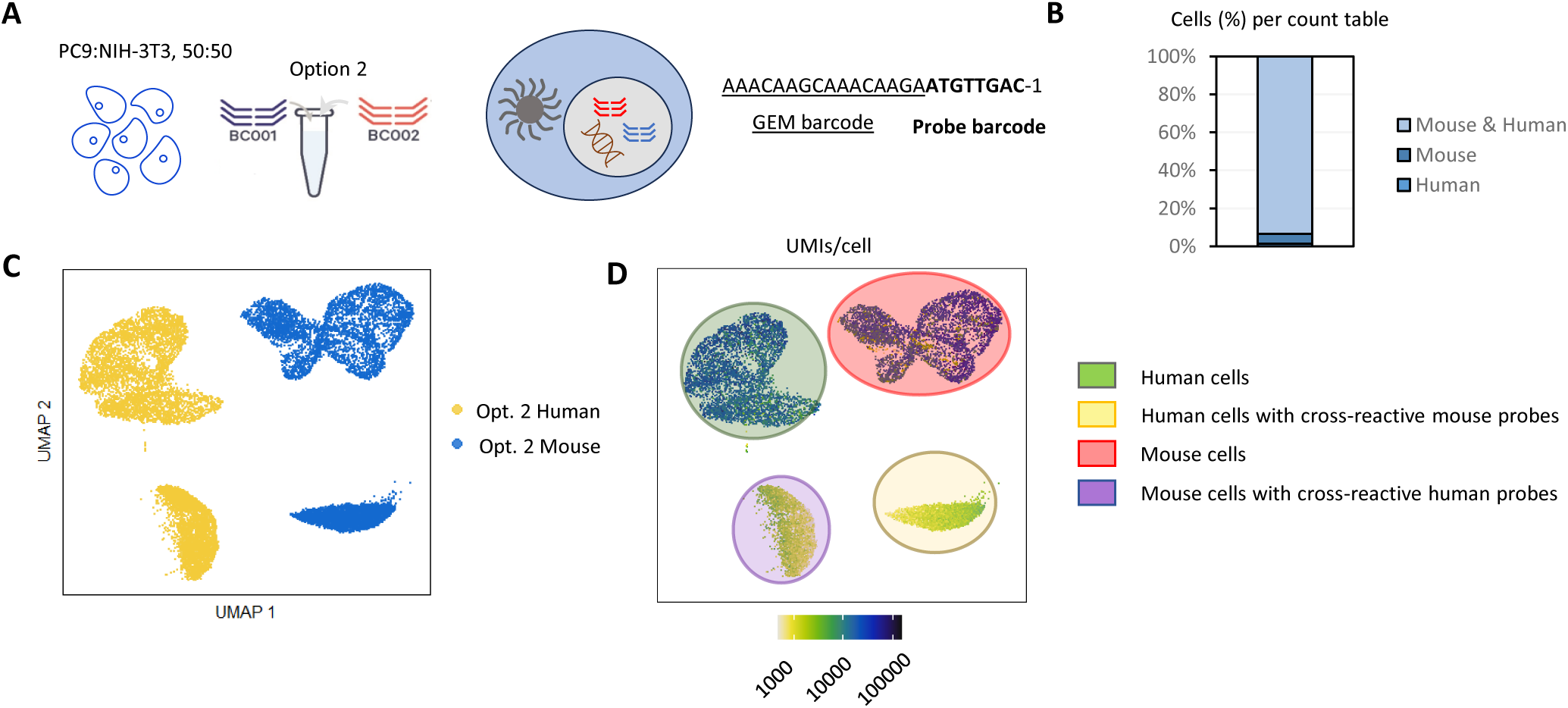
The encapsulation of single cells hybridized to both species probes leads to the presence of the same GEM barcode in each species count table. **A.** Schematic describing the use of a mixed sample with option 2 strategy and the illustration of how the same GEM barcode can end up being present in both count tables. **B.** Bar plot showing the % cells (GEM barcodes) that are present in both Mouse & Human count tables, Mouse count table or Human count table. **C.** UMAP visualization of the clustering of the cells profiled with option 2. Cells profiled with human probes are shown in yellow and the ones profiled with mouse probes are shown in blue. **D.** UMAP visualization with the UMI counts per cell of option 2 sample. Green: human cells. Yellow: human cells profiled by cross-reactive mouse-probes. Red: mouse cells. Purple: mouse cells profiled by cross-reactive human probes.

We next set out to develop a pipeline to correctly classify each cell in each of the count tables. Our assumption is that the true organism probes will have much higher affinity for the transcripts than the cross-hybridized ones. Therefore, our pipeline stringently evaluates the signal in UMI and unique genes for each GEM barcode on each species count table and sorts them accordingly. By doing that, we can classify the cells that are being profiled by cross-hybridized probes for each of the count tables (Fig. 5A). Interestingly the % of cross-reactivity reflects the population structure and the number of true human and mouse cells profiled overall also shows the 50:50 ratio of human and mouse cells present in the sample (Fig. 5B-C). Again, the clustering and UMAPs visualizations confirm that cross-hybridized cells tend to cluster apart from the correctly profiled cells, with lower UMI counts (Fig. 5D-E). In some cases, the classification is unclear because of disagreement between the UMI evaluation and the genes per cell (Fig. 5A). Nevertheless, these are usually low-quality cells that would be anyway filter out during downstream analysis. Our pipeline, therefore, can classify each GEM into the correct species and provide count tables free of cross-reactive signal. It is worth mentioning that the removal of the cross-reactive cells from each count table does not lead to loss of cell numbers as these cells will be correctly profiled in the other count table.

**Figure 5.**
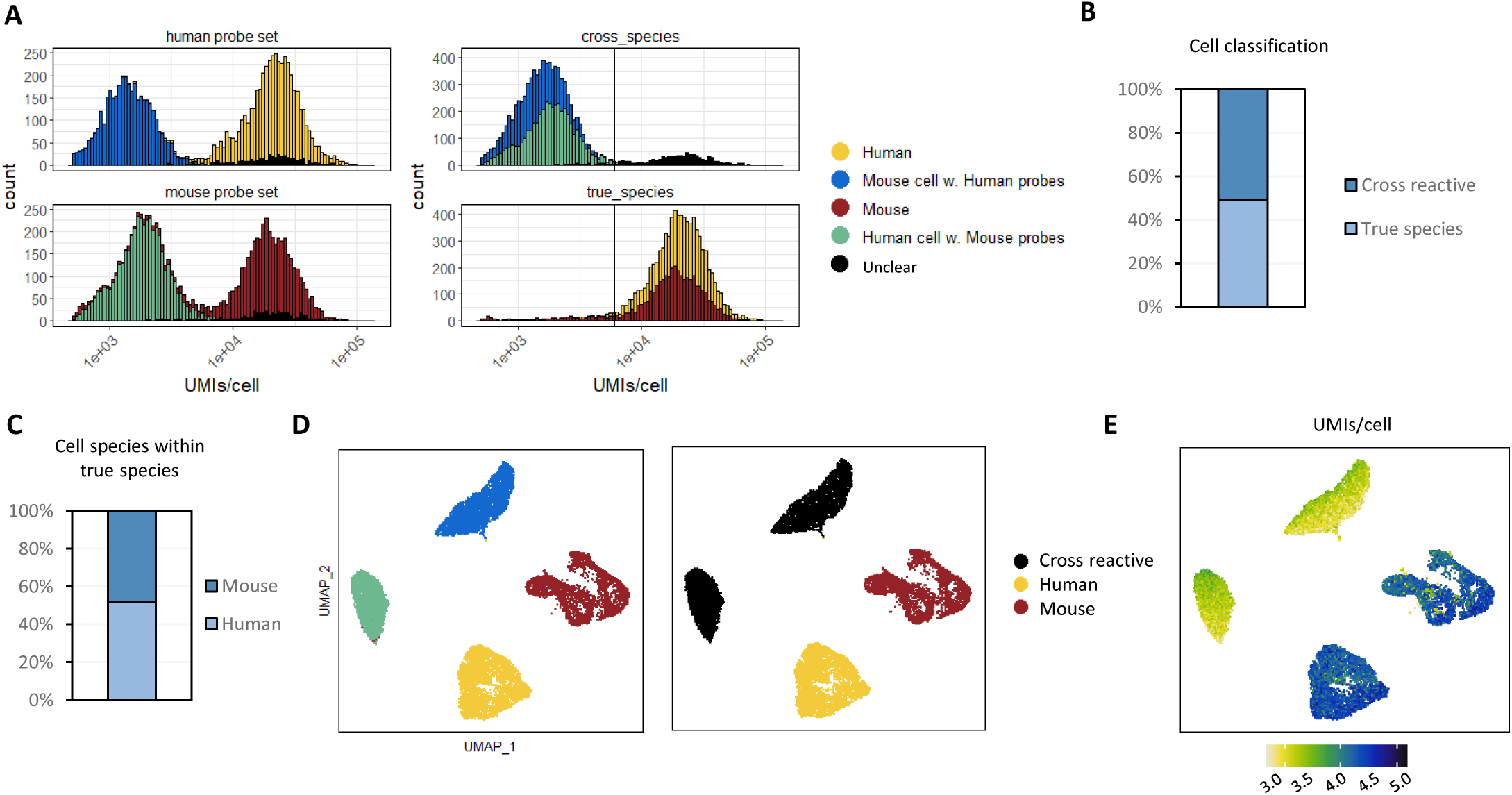
Classification of each GEM barcode into the right species for each of the count tables. **A.** Cumulative cell count histograms showing the distribution of UMIs/cell in the human and mouse count tables after classifying the GEM barcodes into Human (yellow), Mouse cell with Human probes (blue), Mouse (red), Human cell with Mouse probes (olive) or unclear (black). **B.** Bar plot of the % of cells present in both count tables classified as profiled by the cross-reactive probe or the true species probes. **C.** Bar plot of the % of cells classified as Mouse or Human after removing the cross-reactive signal. **D.** UMAP visualizations after clustering of the data within the human and mouse count tables and classifying based on the UMI and unique genes recovered per GEM barcode. **E**. UMAP visualization with the UMI counts per cell.

### Sample sub-pooling and differential loading allow to correct for low input samples

Our next step was to evaluate whether differential pooling of a sub-pool would allow us to recover cells present in lower percentages. This solution would be very helpful in situations where the species of interest represents a low percentage of the initial sample, such as human cells in PDX samples undergoing treatment.

To test this, we used samples with 10% and 25% human cells (PC9), with the rest being mouse cells (NIH-3T3). For each of the samples, we performed a hybridization against human and mouse probes separately (Fig. 6A). Cells were differentially pooled before proceeding to washing and loading into the Chromium X. We aimed to recover similar amounts of cells from both species, thus we adjusted the pooling so that the number of targeted cells would be close to the number recovered in a 50:50 sample. For instance, for the 10:90 sample, we loaded the human hybridization to be a 4.5/8 of the sample and the mouse hybridization was loaded to be a 0.5/8. For the overall pool, we aimed to recover 40,000 cells in total. Therefore, for the 10:90 sample we expected a total recovery of 22,500 cells from the human hybridization and 2,500 cells from the mouse hybridization. Considering the actual percentage of human and mouse cells in the sample, this should produce data for 2,250 human and mouse cells (Fig. 6B).

**Figure 6.**
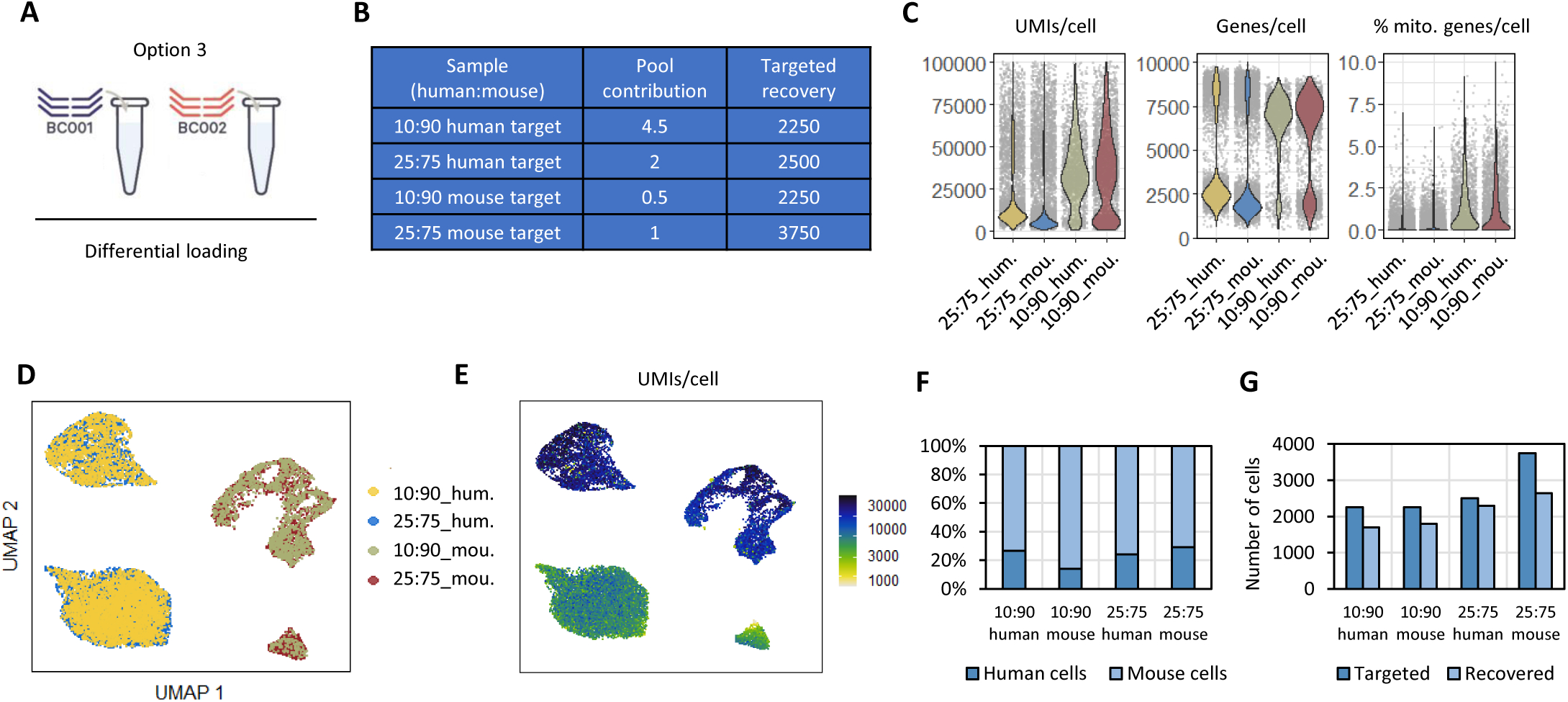
Sub-pooling and differential loading of samples with various percentages of human cells. **A.** Schematic of the sub-pooling strategy for this type of sample: the sample is split into two and hybridized separately to human and mouse probe sets. Then each hybridization can be used into a pool using different amounts of cells to compensate for the low human cell numbers. **B.** Table summarizing the contribution of each sample to the pool and the expected number of cells recovered. **C.** Violin plots showing the distribution of genes, UMI counts and percentage of mitochondrial genes per cell. Yellow: 10:90 sample hybridized to human probes. Blue: 25:75 sample hybridized to human probes. Olive: 10:90 sample hybridized to mouse probes. Red: 25:75 sample hybridized to mouse probes. **D.** UMAP visualization of the clustering obtained from each of the samples. **E.** UMAP visualization with the UMI counts per cell. **F.** Percentage of human and mouse cells per sample when assigning cells present in the low-UMI count clusters as the counterpart species. **G.** Dark blue: number of cells targeted based on the loading of the sub-pool. Light blue: number of cells recovered from the data after filtering out cells within the low UMI count clusters.

When loading the count matrices into a Seurat object we clearly observed the bimodal distribution of UMI counts and genes recovered per cell (Fig. 6C). After processing and clustering we obtained a good overlap between samples and a clear distinction of the cells with low UMI counts, profiled by the cross-reactive probes (Fig. 6D-E). We extracted the number of cells present in the distinct clusters and classified them as human or mouse cells. Cells present in the low UMI counts clusters in the human hybridizations were counted as mouse cells, and vice-versa for the mouse hybridizations. For all the samples, except for the 10:90 human hybridization, the % of human and mouse cells matched the amounts present in the starting material (Fig. 6F). For the 10:90 human hybridization, cellranger set a very conservative threshold on which droplets contain cells, which excludes many cells with low UMI counts from the count matrices. Nevertheless, we recovered the expected number of human cells: we were expecting 2,250 human cells and we recovered 1,700 (Fig. 6G). For the other hybridizations we also recovered most of the expected cells, with a good balance of human and mouse cells targeted by each of the sub-pools (Fig. 6G). Therefore, sample sub-pooling and differential loading allows to compensate uneven species distribution within a sample.

### Profiling of a CDX sample with the 10x Flex kit

Next, we put our pipeline to the test with a PC-9 lung cancer cell-line derived xenograft (CDX) model treated for 7 days with vehicle or an IC90 concentration of an EGFR tyrosine kinase inhibitor (Fig. 7A). We profiled them using option 2 strategy, which demonstrated to be the best approach in the cell line experiments. In this case, around 50% of cells were present in both human and mouse count table (Fig. 7B). The bimodal distribution of the data is also present at the UMI and genes per cell level (Fig. 7C). We applied our classification pipeline and noticed that in the mouse count table, some of these cross-reactive human probes provided levels of UMI counts similar to the ones provided by mouse probes in mouse cells. This highlights the importance of running a more stringent classification like our pipeline does rather than setting UMI or gene thresholds on the bimodal distribution (Fig. 7D). After classifying each GEM into their true species, around 30% of cells in the vehicle sample were mouse cells and we recovered up to 60% in the treated condition (Fig. 7E). The clustering and UMAP visualization also resolved very well the cells with cross-reactive signal with distinctive clusters of cells with low UMI and gene counts (Fig. 7F-G).

**Figure 7.**
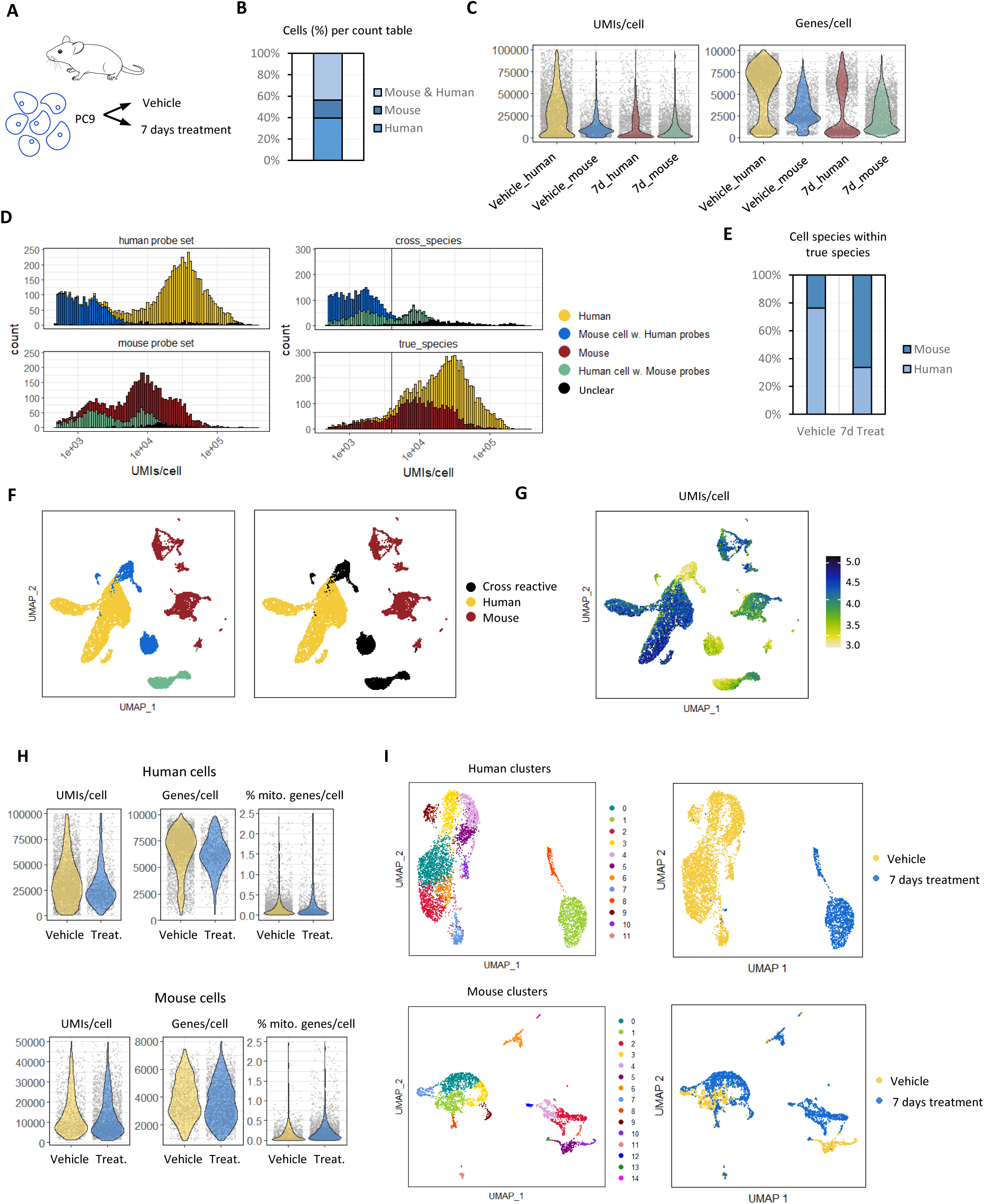
Classifying GEM barcodes into the right species based on UMI and genes per cell in both species count tables allows to perform species specific analysis on CDX samples. **A.** Schematic of the PC9 CDX samples used. **B.** Bar plot showing the % cells (GEM barcodes) that are present in both Mouse & Human count tables, Mouse count table or Human count table. **C.** Violin plots showing the distribution of genes, UMI counts and percentage of mitochondrial genes per cell. Yellow: vehicle sample human count table. Blue: vehicle sample mouse count table. Red: 7 days treated sample human count table. Olive: 7 days treated sample mouse count table. **D.** Cumulative cell count histograms showing the distribution of UMIs/cell after classifying the GEM barcodes into Human (yellow), Mouse cell with Human probes (blue), Mouse (red), Human cell with Mouse probes (olive) or unclear (black). **E.** Bar plot of the % of cells classified as Mouse (dark blue) or Human (light blue) after removing the cross-reactive signal. **F.** UMAP visualizations after clustering of the data within the human and mouse count tables and classifying based on the UMI and unique genes recovered per GEM barcode. **G.** UMAP visualization with the UMI counts per cell. **H.** Violin plots showing the distribution of genes, UMI counts and percentage of mitochondrial genes per cell in the human cells (top) and mouse cells (bottom). **I.** UMAP visualization of the clustering obtained from each of the species (human top, mouse bottom). Yellow: vehicle samples. Blue: 7 days treated samples.

Once the right species is identified for each GEM barcode, we can generate species’ specific count tables, free of cross-reactive signal, which allows downstream analysis for each species separately (Fig. 7H-I).

### Sample sub-pooling in a PDX sample

We used a lung cancer PDX model for which usually around 3% of the cells correspond to the mouse population to test whether we can recover enough cells by differential pooling (Fig. 8A). We tried loading 16 times more volume of the mouse hybridization compared to the human hybridization into the reaction. Therefore, for a target recovery of 30,000 cells we used a 1/17 for the human hybridization leading to an expected recovery of 1,764 cells with 97% being human and a target human cell number of 1,712. For the mouse cells we loaded a 16/17 of the mouse hybridization leading to 28,235 cells with 3% being mouse and a target mouse cell number of 847 cells (Fig. 8B).

**Figure 8.**
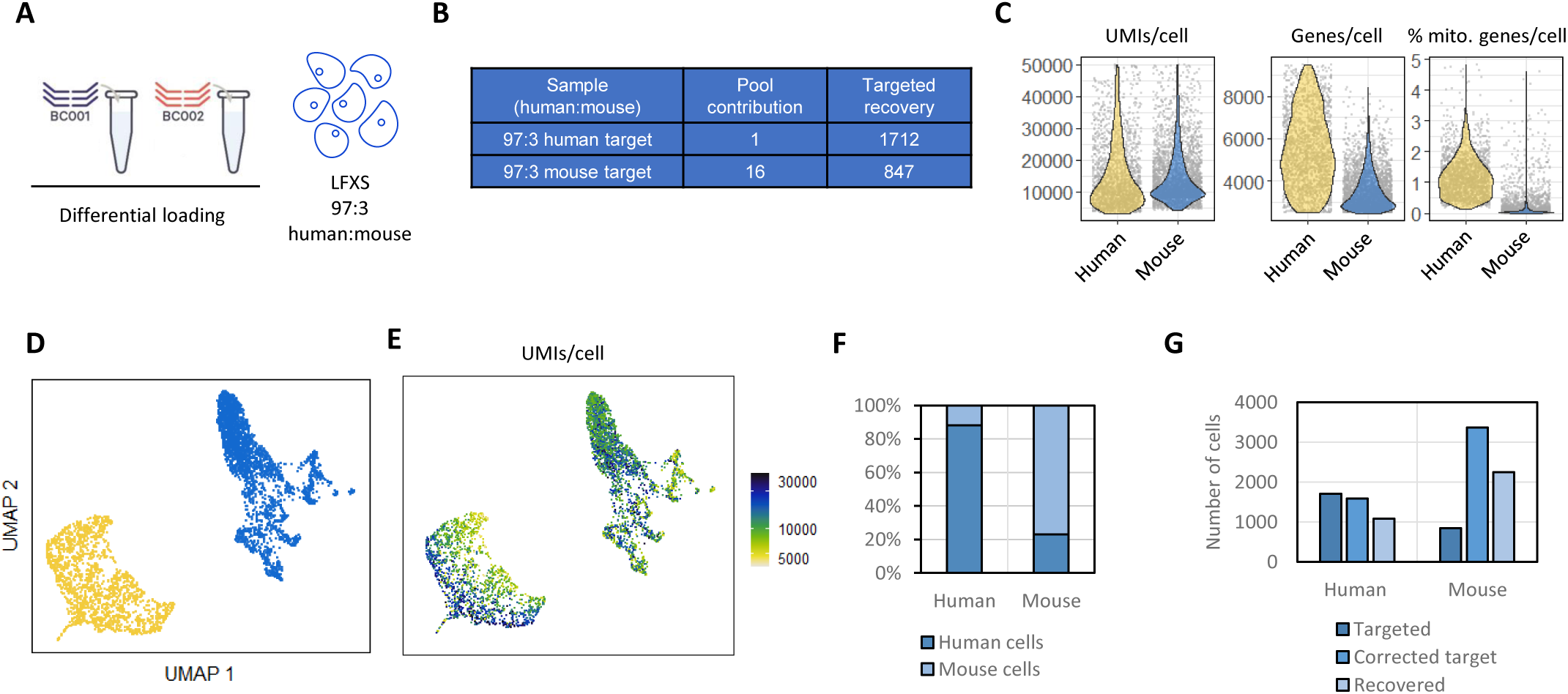
Differential loading of a PDX sample with 3% mouse cells allows to recover enough murine cells. **A.** Schematic of the strategy used to recover a minor murine cell population in a PDX sample. **B.** Table summarizing the contribution of each sample to the pool and the expected number of cells recovered. **C.** Violin plots showing the distribution of genes, UMI counts and percentage of mitochondrial genes per cell. Yellow: human hybridization. Blue: mouse hybridization. **D.** UMAP visualization of the clustering obtained from each of the samples. **E.** UMAP visualization with the UMI counts per cell. **F.** Percentage of human and mouse cells per sample when assigning cells with low-UMI count as the counterpart species. **G.** Bar plot showing the number of cells targeted per hybridization, the number of cells expected to be recovered based on the correction of % of mouse cells observed in the sample and the number of cells recovered.

After running cellranger and exploring the count tables we noticed that again, in the sample being overloaded, many cells got excluded: mostly human cells profiled by cross-reactive probes that did not pass the threshold. Interestingly, when looking at the distribution of genes and UMI per cell the bimodal distribution is not so striking, especially in the human data (Fig. 8C). This is also reflected in the UMAP visualization where the clustering does not allow to identify the cells profiled by cross-reactive probes as easily as with the cell lines (Fig. 8D-E).

Thus, we decided to set a threshold at 3,000 UMI per cell to define the cells profiled by cross-reactive probes. We calculated the number of cells being filtered to calculate the % of human and mouse cells in each of the count tables. For the human count table, 12% of the cells corresponded to mouse, whereas in the mouse count table 80% were mouse cells (Fig. 8F). Again, this reflects the number of cells discarded by cellranger in the mouse count table. The results suggest that the original sample probably contained more mouse cells than the estimated 3%. Therefore, we adjusted the expected cell recovery based on the 12% observed. By doing that, our recovered cell numbers are closer to the expected values (Fig. 8G). Therefore, despite the filtering of cells by cell ranger and the difficulties to establish cross-reactive cells clusters; we are able to discard low quality cells and recover the expected number of human and mouse cells by the differential loading. However, the results suggest that when using a single species probe set per hybridization in PDX samples, it might be difficult to distinguish the cross-reactive signal and therefore the use of both species’ probe sets would be advised.

## DISCUSSION

In vivo xenograft models (whether cell line or patient-derived) are widely used in cancer research to characterise cancer biology, drug resistance and pharmacology. In single-cell sequencing experiments, xenograft samples must typically be dissociated to single-cell and flow sorted to enrich for human cells prior to cryopreservation. These steps may induce transcriptional programs in the sorted cells as well as loss of viable cells. A simplified protocol that does not compromise the quality or depth of single-cell sequencing but avoids these cell sorting steps would have logistic, biological and financial advantages.

We therefore sought to test the feasibility of using a novel application of the existing Gene Expression Flex protocol solution that combines the human and mouse gene probes in the same sample.

Our results point to cross-reactivity levels being sufficiently low to correctly identify the correct species for each of the cells present in a sample. By mixing the human and mouse probe sets within the same hybridization, we can profile both populations while discarding the signal provided by the cross-reactive probes. Cellranger facilitates this classification by providing a count table for each of the barcodes used in a particular sample, in this case, a count table from the human probes and a count table from the mouse probes. The presence of cross-reactive probes has minimal effects on the gene expression profiles obtained. Most of the genes showing gene expression differences are targeted by single probes and are highly conserved, therefore might suffer from competition in presence of both human and mouse transcripts. Additionally, the processing of a sample into two different species hybridizations can be exploited to increase the recovery of a particular species population in cases where it is expected to be underrepresented.

In conclusion, we demonstrate the feasibility of profiling PDX and CDX models with the Gene Expression Flex kit by mixing the species’ probes sets and without any appreciable loss in sequencing quality. We anticipate that in future fixation protocols which minimize the handling of single-cells from in *vivo* models will greatly simplify sample collection as well as the ability to interrogate simultaneously the human and mouse transcriptome from such samples.

## MATERIALS AND METHODS

### Biological materials

This study was carried out in strict accordance with the recommendations in the Guide for the Care and Use of Laboratory Animals of the Society of Laboratory Animals (GV SOLAS) in an AAALAC accredited animal facility. All animal experiments were approved by the Committee on the Ethics of Animal Experiments of the regional council (Permit Numbers: I-19/02). Non-small cell lung cancer (NSCLC) PDX models were implanted subcutaneously in 4– 6-week-old female NMRI nude mice (Charles River, Germany) under isoflurane anesthesia. Tumor growth was determined by a two-dimensional measurement with calipers twice a week. Different analyses of the PDX models were performed when tumor size reached 400 – 500 mm³. Animals were sacrificed and tumors were sampled for subsequent analysis.

PC9 CDX animal studies were conducted in accordance with U.K. Home Office legislation, the Animal Scientific Procedures Act 1986, and AstraZeneca Global Bioethics policy. All experimental work is outlined in project license PP3292652, which has gone through the AstraZeneca Ethical Review Process. Female severe combined immunodeficiency disease (SCID) mice were purchased from Envigo UK. All mice were older than 6 weeks at the time of implant. Cells were implanted under isoflurane anaesthesia with microchipping performed at same time. Approximately 5 million PC9 cells in a total volume of 0.1 mL containing 50% Matrigel were injected subcutaneously in the right flank of female mice. PC9 xenografts were established in female SCID mice. Mice were marked with a subcutaneous microchip to follow tumour growth and tumours were measured with an electronic calliper twice weekly. Tumour volume was calculated as 0.5 x tumour length x tumour width x 2. Tumour volume was monitored twice weekly by bilateral calliper measurements until the end of study or until tumours reached the ethical limit of 10% of the mouse’s bodyweight.

### Sample preparation: fixation and storage

PC9 cells were grown in RPMI 1640 with Glutamax (Gibco) and 10% FBS, while NIH-3T3 cells were grown in DMEM (Gibco) with L-glutamine and 10% FBS. Cells were dissociated using Tryple Express (no phenol red) (Gibco) and made into single-cell suspension, followed by filtration through 30uM Miltenyi pre-separation filters and counting using AO/PI double stain (Nexcelom). Cells were processed according to the 10x protocol for the fixation of cells and nuclei. Briefly, suspensions of up to 10 million cells were spun-down at 350 rcf for 5 min at 4C and cells were resuspended in 1 mL of Fixation Buffer (4% Formaldehyde, 1X Conc. Fix and Perm Buffer – 10x Genomics PN-2000517). Cells were fixed for 16-24h at 4C. To stop the fixation, cells were spun-down at 850 rcf for 5 min at room temperature and quenched with 1 mL of Quenching Buffer (1X Conc. Quench Buffer – 10x Genomics PN-2000516). Preparations were then processed by adding 0.1 volumes of Enhancer (10x Genomics PN-2000482) and 10% glycerol for long-term storage and shipment to Single-cell Discoveries B.V.. Samples were then thawed at room temperature, centrifuged at 850 rcf for 5 min and resuspended in 1 mL 0.5X PBS - 0,02% BSA. Cell concentration and viability was then assessed by AO-PI staining with a LUNA-FX7™ Automated Cell Counter (Logos Biosystems).

Mice were randomized into vehicle or treatment groups with approximate mean tumour volume of 0.2 to 0.4 cm3. Randomization for animal studies was based on initial tumour volumes to ensure equal distribution across groups. Mice were dosed once in the morning daily by oral gavage for the duration of the treatment period. All compounds were synthesised by AstraZeneca. TKI (osimertinib) was prepared fresh every 7 days in 0.5% HPMC. Mice were sacrificed at indicated time point and the collected tumours were passed through a McIlwain tissue chopper and stored in 1mL RPMI1640 until subsequent processing. Minced PC9 CDX tissue samples were dissociated using the Miltenyi human tumour dissociation kit (130-095-929) (40 minutes, 37C), filtered through 30uM Miltenyi pre-separation filters, counted using AO/PI double stain (Nexcelom) and processed as per 10x Flex protocol for fixation of dissociated tumour cells.

### Sample hybridization

Up to 2 million cells were processed per hybridization following 10x recommendations. Hybridizations were set up in 80 uL of hybridization mix with 20 uL of Human or Mouse WTA probes (10x Genomics PN-2000510 or PN-2000718). For experiments with probe mixing 10 uL of each probe set were used per hybridization. Hybridizations were performed at 42C for 16-24h. After hybridization samples were diluted in Post-Hyb Wash Buffer and measured by AO-PI staining in a LUNA-FX7™ Automated Cell Counter (Logos Biosystems). For each experiment, we pooled an equal number of cells from each hybridization to have an equal contribution per sample. For the experiments using sub-pools and differential loading, cell numbers were adjusted to the desired contribution in the pool as indicated. Cell pools were then washed 3 times in Post-Hyb Wash Buffer for 10 min at 42C. After the washes cells were resuspended in Post-Hyb Resuspension Buffer, filtered through a Miltenyi Biotec 30 um filter and measured with the cell counter to determine the amount needed for the Chromium X run.

### Gene Expression Flex run

For the GEM encapsulation we followed the 10x Genomics protocol and their guidelines on the volume of cells and reagents required per well according to the targeted cell recovery. For the PC9 only, spiked samples and PDXs we targeted 8,000 cells per sample. For the experiment testing probe mixing strategies, we targeted 40,000 cells per pool, with equal contribution per sample except for the differential pooling strategy where each sample contributed differently. After loading the Chip Q and running it on the Chromium X, GEMs were recovered and processed as indicated by 10x Genomics. After processing the GEMs, the product is pre-amplified and indexed to construct the sequencing library.

### Sequencing

Libraries were sent for sequencing to the Harwig Medical Foundation with the Illumina Nova-Seq 6000 or internally at Single-cell Discoveries with the Illumina Nova-Seq X sequencing platform following 10x recommended settings. For the PC9 spiked samples we sequenced at a depth of 15,000 reads. For the experiments with the probe mixing strategies and PDXs, we sequenced at a depth of 30,000 reads per cell.

### Cellranger multi analysis

10x Genomics cellranger v7.1.0 was used for all analyses. A combined human and mouse genome reference was constructed by concatenating the human and mouse FASTA and GTF files provided by 10x Genomics, respectively. The human reference used was GRCh38-2020-A (GRCh38, GENCODE version 32 (Ensembl 98), with 10x Genomics modifications, see https://support.10xgenomics.com/single-cell-gene-expression/software/release-notes/build#GRCh38_2020A). The mouse reference used was mm10-2020-A (GRCm38, version GENCODE M23 (Ensembl 98), with 10x Genomics modifications, see https://support.10xgenomics.com/single-cell-gene-expression/software/release-notes/build#mm10_2020A). A cellranger reference was then created using the combined FASTA and GTF files using the ‘cellranger mkref’ command with default parameters. Raw 10x Flex data (FASTQ) was mapped to the combined human+mouse reference using ‘cellranger multi’ command.

### Single-cell data analysis

Count matrices were used to create a Seurat (v4.3) object excluding cells with less than 200 genes and genes present in less than 3 cells. Data was log-normalized and the top 2,000 variable genes were selected to guide the dimensionality reduction by PCA. Data was scaled and clustering was performed following the Seurat package with a cluster resolution of 0.5. The manifold was then visualized with a UMAP representation. Clusters suspected to be human and mouse were subset and differential expression analysis was performed using a Wilcoxon Rank Sum test.

## LIMITATIONS OF THE STUDY

The use of the probe sets limits the analysis to roughly 18,000-19,000 protein coding genes, excluding TCR, IG, ribosomal protein, mitochondrial ribosomal genes, KIR, HLA and non-coding RNA. The cross-reactive signal will likely be sample-specific, and it is difficult to quantify the impact on the gene expression levels captured by the specific probe binding. From the results obtained so far it seems that most of the differences are mainly observed in genes covered by a single probe and other few genes with high homology between the two species. In any case, most studies will include a set of control and test samples that should suffer from the same cross-reactivity, which will not impact the hypothesis testing when comparing control and tests. From our results we would advise to always include both human and mouse probe sets to profile PDX samples, which would simplify the classification of true signal and cross-reactive signal.

## CONTRIBUTIONS

O. Llorà-Batlle, A. Farcas, K. Lashuk, J. Schueler, D. Mooijman and U. McDermott designed the experiments. O. Llorà-Batlle, A. Farcas and K. Lashuk performed the experiments. A. Farcas, N. Floc’h, S. Talbot, A. Schwiening, L. Bojko and J. Calver were in charge of the AZ mouse models and prepared the corresponding samples. O. Llorà-Batlle and A. Prodan performed data analysis. O. Llorà-Batlle, A. Farcas, D. Mooijman and U. McDermott wrote the manuscript. The PERSIST-SEQ project has received funding from the Innovative Medicines Initiative 2 www.imi.europa.eu Joint Undertaking under grant agreement No 101007937. This Joint Undertaking receives support from the European Union’s Horizon 2020 research and innovation program and EFPIA.

## FIGURES

**Supplementary Figure 1.**
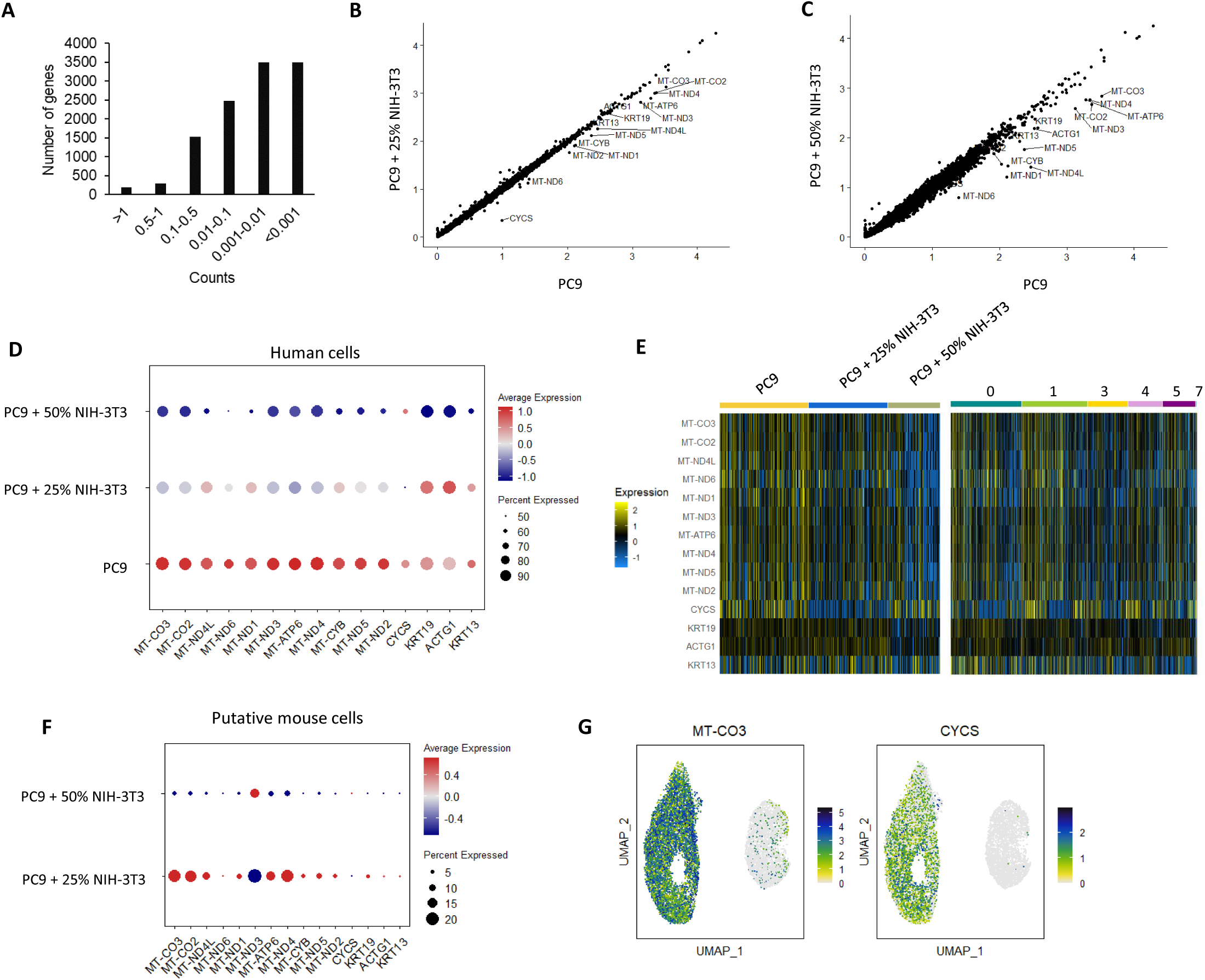
QC on the difference between the PC9 cells recovered from the un-spiked and spiked samples. **A.** Distribution of average UMI counts per gene in the cells within the cross-reactive clusters. **B.** Correlation between the average gene counts of the PC9 cells against the PC9 cells in the 25% spiked sample. **C.** Correlation between the average gene counts of the PC9 cells against the PC9 cells in the 50% spiked sample. **D.** Dot plot showing the average expression level of the genes differentially expressed between the human cells in the un-spiked and spiked samples. Expression levels are shown by the blue to red scale and the percentage of cells expressing the gene is shown by the radius of the dot. **E.** Heatmap displaying the differentially expressed genes between human cells in the un-spiked and spiked samples plotted by sample (left) and by cluster (right). **F.** Dot plot showing the average expression level of the genes differentially expressed between human cells in the putative mouse clusters. **G.** UMAP visualizations of the normalized expression levels of MT-CO3 and CYCS.

**Supplementary Figure 2.**
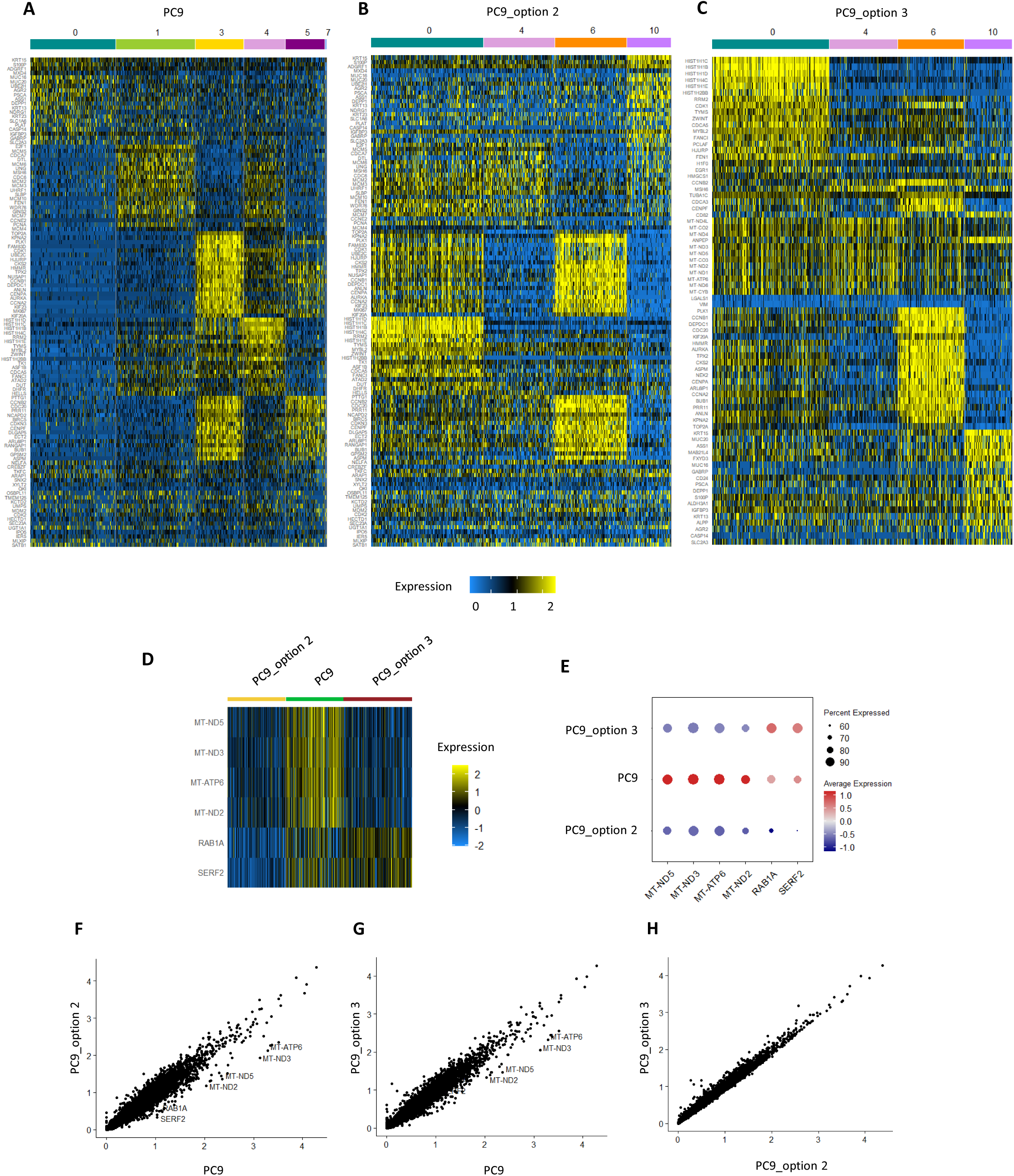
PC9 cells profiled with the different strategies are very similar. **A.** Heatmap showing the top 10 marker genes per cluster in PC9 cells. Minimum expression in 50% of cells and log2FC>0.25. **B.** Heatmap showing the top 10 marker genes per cluster in PC9 cells profiled by mixing probes with different barcodes (option 2). **C.** Heatmap showing the top 10 marker genes per cluster in PC9 cells profiled by sub-pooling and hybridizing to human probes (option 3). **D.** Heatmap showing the differentially expressed genes between the PC9 cells for each of the samples. Minimum expression in 25% of cells and log2FC>0.5. **E.** Dot plot showing the average expression level of the genes differentially expressed between PC9 cells in each of the samples. Expression levels are shown by the blue to red scale and the percentage of cells expressing the gene is shown by the radius of the dot. **F.** Correlation between the average gene counts of PC9 cells against the PC9 cells from the mixed sample profiled with option 2 strategy. **G.** Correlation between the average gene counts of PC9 cells against the PC9 cells from the mixed sample profiled with option 3 strategy. **H.** Correlation between the average gene counts of PC9 cells profiled with option 2 and option 3 strategies.

